# The evolution, diversity and host associations of rhabdoviruses

**DOI:** 10.1101/020107

**Authors:** Ben Longdon, Gemma GR Murray, William J Palmer, Jonathan P Day, Darren J Parker, John J Welch, Darren J Obbard, Francis M Jiggins

**Affiliations:** Department of Genetics University of Cambridge Cambridge CB2 3EH UK; School of Biology University of St. Andrews St. Andrews KY19 9ST UK; Department of Biological and Environmental Science, University of Jyväskylä, Jyväskylä, Finland; Institute of Evolutionary Biology, and Centre for Immunity Infection and Evolution University of Edinburgh Edinburgh EH9 3JT UK

**Keywords:** Virus, Host shift, Arthropod, Insect, Rhabdoviridae, Mononegavirales

## Abstract

Metagenomic studies are leading to the discovery of a hidden diversity of RNA viruses. These new viruses are poorly characterised and new approaches are needed predict the host species these viruses pose a risk to. The rhabdoviruses are a diverse family of RNA viruses that includes important pathogens of humans, animals and plants. We have discovered 32 new rhabdoviruses through a combination of our own RNA sequencing of insects and searching public sequence databases. Combining these with previously known sequences we reconstructed the phylogeny of 195 rhabdovirus sequences, and produced the most in depth analysis of the family to date. In most cases we know nothing about the biology of the viruses beyond the host they were identified from, but our dataset provides a powerful phylogenetic approach to predict which are vector-borne viruses and which are specific to vertebrates or arthropods. By reconstructing ancestral and present host states we found that switches between major groups of hosts have occurred rarely during rhabdovirus evolution. This allowed us to propose 76 new likely vector-borne vertebrate viruses among viruses identified from vertebrates or biting insects. Based on currently available data, our analysis suggests it is likely there was a single origin of the known plant viruses and arthropod-borne vertebrate viruses, while vertebrate-specific and arthropod-specific viruses arose at least twice. There are also few transitions between aquatic and terrestrial ecosystems. Viruses also cluster together at a finer scale, with closely related viruses tending to be found in closely related hosts. Our data therefore suggest that throughout their evolution, rhabdoviruses have occasionally jumped between distantly related host species before spreading through related hosts in the same environment. This approach offers a way to predict the most probable biology and key traits of newly discovered viruses.

## Introduction

RNA viruses are an abundant and diverse group of pathogens. In the past, viruses were typically isolated from hosts displaying symptoms of infection, before being characterized morphologically and then sequenced following PCR [1, 2]. PCR-based detection of novel RNA viruses is problematic as there is no single conserved region of the genome shared by all viruses from a single family, let alone across all RNA viruses. High throughput next generation sequencing technology has revolutionized virus discovery, allowing rapid detection and sequencing of divergent virus sequences simply by sequencing total RNA from infected individuals [1, 2]

One particularly diverse family of RNA viruses is the *Rhabdoviridae.* Rhabdoviruses are negative-sense single-stranded RNA viruses in the order *Mononegavirales* [3]. They infect an extremely broad range of hosts and have been discovered in plants, fish, mammals, reptiles and a broad range of insects and other arthropods [4]. The family includes important pathogens of humans and livestock. Perhaps the most well-known is rabies virus, which can infect a diverse array of mammals and causes a fatal infection killing 59,000 people per year with an estimated economic cost of $8.6 billion (US) [5]. Other rhabdoviruses, such as vesicular stomatitis virus and bovine ephemeral fever virus, are important pathogens of domesticated animals, while others are pathogens of crops [3].

Arthropods play a key role in the transmission of many rhabdoviruses. Many viruses found in vertebrates have also been detected in arthropods, including sandflies, mosquitoes, ticks and midges [6]. The rhabdoviruses that infect plants are also often transmitted by arthropods [7] and some that infect fish can potentially be vectored by ectoparasitic copepod sea-lice [8, 9]. Moreover, insects are biological vectors; rhabdoviruses replicate upon infection of insect vectors [7]. Other rhabdoviruses are insect-specific. In particular, the sigma viruses are a clade of vertically transmitted viruses that infect dipterans and are well-studied in *Drosophila* [10-12]. Recently, a number of rhabdoviruses have been found to be associated with a wide array of insect and other arthropod species, suggesting they may be common arthropod viruses [13, 14]. Furthermore, a number of arthropod genomes contain integrated endogenous viral elements (EVEs) with similarity to rhabdoviruses, suggesting that these species have been infected with rhabdoviruses at some point in their history [15-18].

Here we explore the diversity of the rhabdoviruses, and examine how they have switched between different host taxa during their evolutionary history. Insects infected with rhabdoviruses commonly become paralysed on exposure to CO_2_ [19-21]. We exploited this fact to screen field collections of flies from several continents for novel rhabdoviruses that were then sequenced using metagenomic RNA-sequencing (RNA-seq). Additionally we searched for rhabdovirus-like sequences in publicly available RNA-seq data. We identified 32 novel rhabdovirus-like sequences from a wide array of invertebrates and plants, and combined them with recently discovered viruses to produce the most comprehensive phylogeny of the rhabdoviruses to date. For many of the viruses we do not know their true host range, so we used the phylogeny to identify a large number of new likely vector-borne viruses and to reconstruct the evolutionary history of this diverse group of viruses.

## Methods

### Discovery of new rhabdoviruses by RNA sequencing

Diptera (flies, mostly Drosophilidae) were collected in the field from Spain, USA, Kenya, France, Ghana and the UK (Data S1: http://dx.doi.org/10.6084/m9.figshare.1425432]. Infection with rhabdoviruses can cause *Drosophila* and other insects to become paralysed after exposure to CO_2_ [19-21], so we enriched our sample for infected individuals by exposing them to CO_2_ at 12°C for 15 mins, only retaining individuals that showed symptoms of paralysis 30mins later. We extracted RNA from 79 individual insects (details in Data S1 http://dx.doi.org/10.6084/m9.figshare.1425432] using Trizol reagent (Invitrogen) and combined the extracts into two pools (retaining non-pooled individual RNA samples). RNA was then rRNA depleted with the Ribo-Zero Gold kit (epicenter, USA) and used to construct Truseq total RNA libraries (Illumina). Libraries were constructed and sequenced by BGI (Hong Kong) on an Illumina Hi-Seq 2500 (one lane, 100bp paired end reads, generating ~175 million reads). Sequences were quality trimmed with Trimmomatic (v3); Illumina adapters were clipped, bases were removed from the beginning and end of reads if quality dropped below a threshold, sequences were trimmed if the average quality within a window fell below a threshold and reads less than 20 base pairs in length were removed. We *de novo* assembled the RNA-seq reads with Trinity (release 2013-02-25) using default settings and jaccard clip option for high gene density. The assembly was then searched using tblastn to identify rhabdovirus-like sequences, with known rhabdovirus coding sequences as the query. Any contigs with high sequence similarity to rhabdoviruses were then reciprocally compared to Genbank cDNA and RefSeq nucleotide databases using tblastn and only retained if they most closely matched a virus-like sequence. Raw read data were deposited in the NCBI Sequence Read Archive (SRP057824). Putative viral sequences have been submitted to Genbank (accession numbers in Tables S1 and S2).

As the RNA-seq was performed on pooled samples, we assigned rhabdovirus sequences to individual insects by PCR on RNA from individual samples. cDNA was produced using Promega GoScript Reverse Transcriptase and random-hexamer primers, and PCR performed using primers designed from the rhabdovirus sequences. Infected host species were identified by sequencing the mitochondrial gene *COI.* We were unable to identify the host species of the virus from a *Drosophila affinis* sub-group species (sequences appear similar to both *Drosophila affinis* and the closely related Drosophila *athabasca*), despite the addition of further mitochondrial and nuclear sequences to increase confidence. In all cases we confirmed that viruses were only present in cDNA and not in non reverse-transcription (RT) controls (i.e. DNA) by PCR, and so they cannot be integrated into the insect genome (i.e. endogenous virus elements or EVEs [17]). *COI* primers were used as a positive control for the presence of DNA in the non RT template. We identified sigma virus sequences in RNA-seq data from *Drosophila montana* [22]. We used RT-PCR on an infected fly line to amplify the virus sequence, and carried out additional Sanger sequencing with primers designed using the RNA-seq assembly. Additional virus sequences were identified from an RNA-seq analysis of pools of wild caught *Drosophila:* DImmSV from *Drosophila immigrans* (collection and sequencing described [23]), DTriSV from a pool of *Drosophila tristis* and SDefSV from *Scaptodrosophila deflexa* (both Darren Obbard, unpublished data. Genbank accession numbers for new virus sequences are (KR822817, KR822816, KR822823, KR822813, KR822820, KR822821, KR822822, KR822815, KR822824, KR822812, KR822811, KR822814 and KR822818). A full list of accessions can be found in tables S1 and S2.

### Discovery of rhabdoviruses in public sequence databases

Rhabdovirus L gene sequences were used as queries to search (tblastn) expressed sequence tag (EST) and transcriptome shotgun assembly (TSA) databases (NCBI). All sequences were reciprocally BLAST searched against Genbank cDNA and RefSeq databases and only retained if they matched a virus-like sequence. We used two approaches to examine whether sequences were present as RNA but not DNA. First, where assemblies of whole-genome shotgun sequences were available, we used BLAST to test whether sequences were integrated into the host genome. Second, for the virus sequences in the butterfly *Pararge aegeria* and the medfly *Ceratitis capitata* we were able to obtain infected samples to confirm whether sequences are only present in RNA by performing PCR on both genomic DNA and cDNA as described above (samples kindly provided by Casper Breuker/Melanie Gibbs, and Philip Leftwich respectively)

### Phylogenetic analysis

All available rhabdovirus-like sequences were downloaded from Genbank (accessions in Data S2: http://dx.doi.org/10.6084/m9.figshare.1425419). Amino acid sequences for the L gene (encoding the RNA Dependent RNA Polymerase or RDRP) were used to infer the phylogeny (L gene sequences: http://dx.doi.org/10.6084/m9.figshare.1425067), as they contain conserved domains that can be aligned across this diverse group of viruses. Sequences were aligned with MAFFT [24] under default settings and then poorly aligned and divergent sites were removed with either TrimAl (v1.3 strict settings, implemented on Phylemon v2.0 server, alignment: http://dx.doi.org/10.6084/m9.figshare.1425069) [25] or Gblocks (v0.91b selecting smaller final blocks, allowing gap positions and less strict flanking positions to produce a less stringent selection, alignment: http://dx.doi.org/10.6084/m9.figshare.1425068) [26]. These resulted in alignments of 1492 and 829 amino acids respectively.

Phylogenetic trees were inferred using Maximum Likelihood in PhyML (v3.0) [27] using the LG substitution model [28] (preliminary analysis confirmed the results were robust to the amino acid substitution model selected), with a gamma distribution of rate variation with four categories and a sub-tree pruning and regrafting topology searching algorithm. Branch support was estimated using Approximate Likelihood-Ratio Tests (aLRT) that are reported to outperform bootstrap methods [29]. Figures were created using FIGTREE (v. 1.4) [30].

### Analysis of phylogenetic structure between viruses taken from different hosts and ecologies

We measured the degree of phylogenetic structure between virus sequences identified in different categories of host (arthropods, vertebrates and plants) and ecosystems (terrestrial and aquatic). Following Bhatia et al [32], we measured the degree of genetic structure between virus sequences from different groups of hosts/ecosystems using Hudson’s *F_st_* estimator [31] as in [32]. We calculated *F_st_* as: 1-the mean number of differences between sequences within or between populations, where a population is a host category or ecosystem. The significance of this value was tested by comparison with 1000 replicates with host categories randomly permuted over sequences. We also measured the clustering of these categories over our phylogeny using the genealogical sorting index (GSI), a measure of the degree of exclusive ancestry of a group on a rooted genealogy [33], for each of our host association categories. The index was estimated using the genealogicalSorting R package [34], with significance estimated by permutation. The tree was pruned to remove strains that could not be assigned to one of the host association categories under consideration. Finally, since arthropods are the most sampled host, we tested for evidence of genetic structure within the arthropod-associated viruses that would suggest co-divergence with their hosts or preferential host-switching between closely related hosts. We calculated the Pearson correlation coefficient of the evolutionary distances between viruses and the evolutionary distances between their hosts and tested for significance by permutation (as in [35]). We used the patristic distances of our ML tree for the virus data and a time-tree of arthropod genera, using published estimates of divergence dates [36, 37].

### Reconstruction of host associations

Viruses were categorised as having one of four types of host association: arthropod-specific, vertebrate-specific, arthropod-vectored plant, or arthropod-vectored vertebrate. However, the host association of some viruses are uncertain when they have been isolated from vertebrates, biting-arthropods or plant-sap-feeding arthropods. Due to limited sampling it was not clear whether viruses isolated from vertebrates were vertebrate specific or arthropod-vectored vertebrate viruses; or whether viruses isolated from biting-arthropods were arthropod specific viruses or arthropod-vectored vertebrate viruses; or if viruses isolated from plant-sap-feeding arthropods were arthropod-specific or arthropod-vectored plant viruses.

We classified a virus from a nematode as having its own host category. We classified three of the fish infecting dimarhabdoviruses as vertebrate specific based on the fact they can be transmitted via immersion in water containing virus during experimental conditions [38-40], and the widely held belief amongst the fisheries community that these viruses are not typically vectored [8]. However, there is some evidence these viruses can be transmitted by arthropods (sea lice) in experiments [8, 9] and so we would recommend this be interpreted with some caution. Additionally, although we classified the viruses identified in sea-lice as having biting arthropod hosts, they may be crustacean-specific. The two viruses from *Lepeophtheirus salmonis* do not seem to infect the fish they parasitise and are present in all developmental stages of the lice, suggesting they may be transmitted vertically [41].

We simultaneously estimated both the current and ancestral host associations, and the phylogeny of the viruses, using a Bayesian analysis, implemented in BEAST v1.8 [42, 43]. Since meaningful branch lengths are essential for this analysis (uncertainty about branch lengths will feed into uncertainty about the estimates), we used a subset of the sites and strains used in the Maximum Likelihood (ML) analysis. We retained 189 taxa; all rhabdoviruses excluding the divergent fish-infecting novirhabdovirus clade and the virus from *Hydra,* as well as the viruses from *Lolium perenne and Conwentzia psociformis,* which had a large number of missing sites. Sequences were trimmed to a conserved region of 414 amino acids where data was recorded for most of these viruses (the Gblocks alignment trimmed further by eye: http://dx.doi.org/10.6084/m9.figshare.1425431).

We used the host-association categories described above, which included ambiguous states. To describe amino acid evolution we used an LG substitution model with gamma distributed rate variation across sites [28] and an uncorrelated lognormal relaxed clock model of rate variation among lineages [44]. To describe the evolution of the host associations we used a strict clock model and a discrete asymmetric transition rate matrix (allowing transitions to and from a host association to take place at different rates), as previously used to model migrations between discrete geographic locations [45] and host switches [43, 46]. We also examined how often these viruses jumped between different classes of hosts using reconstructed counts of biologically feasible changes of host association and their HPD confidence intervals (CI) using Markov Jumps [47]. These included switches between arthropod-specific and both arthropod-vectored vertebrate and arthropod-vectored plant states, and between vertebrate specific and arthropod-vectored vertebrate states. We used a constant population size coalescent prior for the relative node ages (using a birth-death prior gave equivalent results) and the BEAUti v1.8 default priors for all other parameters [42] (BEAUti xml http://dx.doi.org/10.6084/m9.figshare.1431922). In Figure 2 we have transferred the ancestral state reconstruction from the BEAST tree to the maximum likelihood tree.

**Figure 1.**
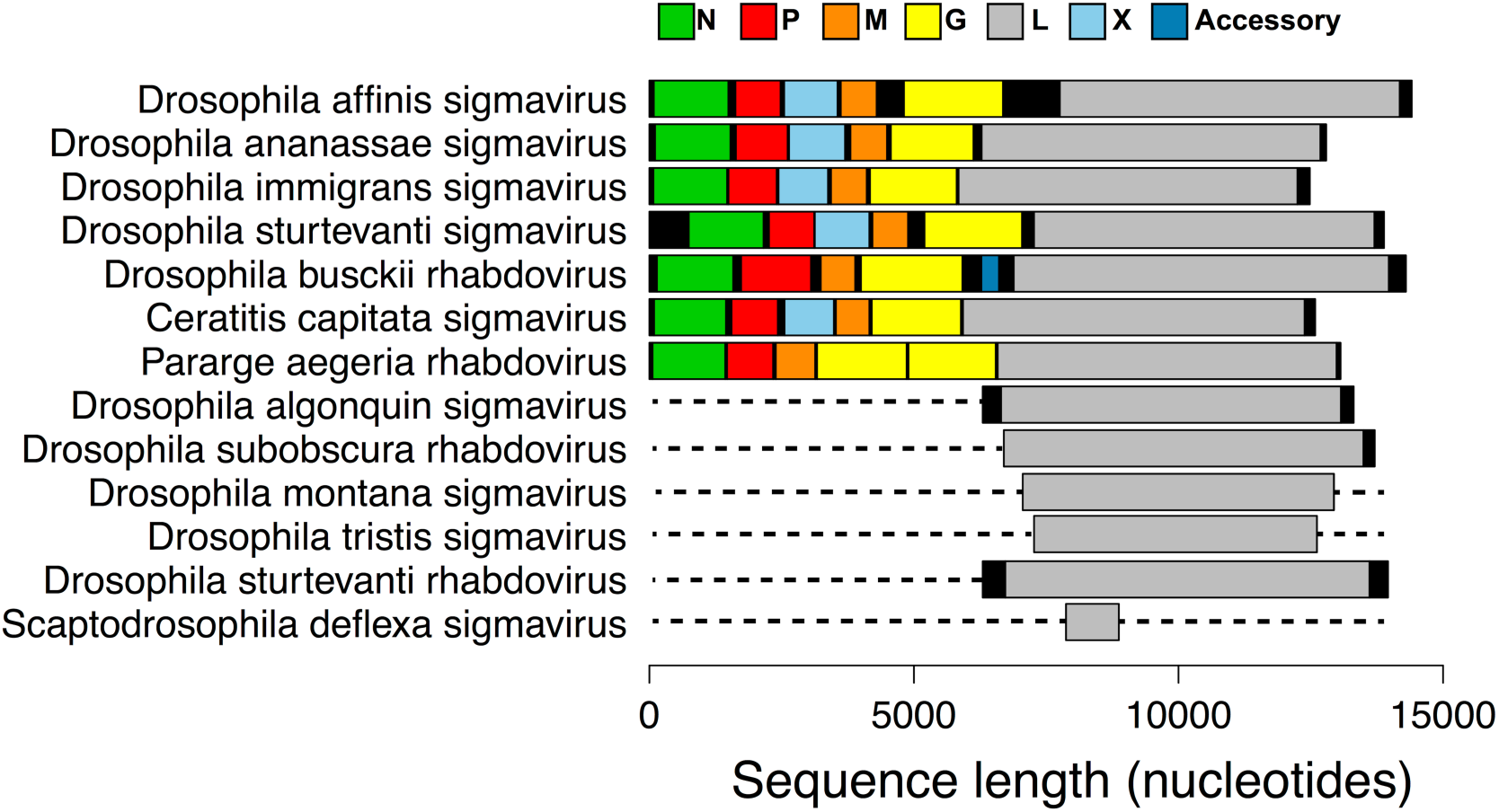
Genome organization of newly discovered viruses from metagenomic RNA sequencing of CO_2_ sensitive flies. Putative genes are shown in colour, non-coding regions are shown in black. ORFs were designated as the first start codon following the transcription termination sequence (7 U’s) of the previous ORF to the first stop codon. Dotted lines represent parts of the genome not sequenced. These viruses were either from our own RNA-seq data, or were first found in in public databases and key features verified by PCR and Sanger sequencing. Rhabdovirus genomes are typically ~11-13kb long and contain five core genes 3’-N-P-M-G-L-5’ [3]. However, a number of groups of rhabdoviruses contain additional accessory genes and can be up to ~16kb long [14, 52].

**Figure 2.**
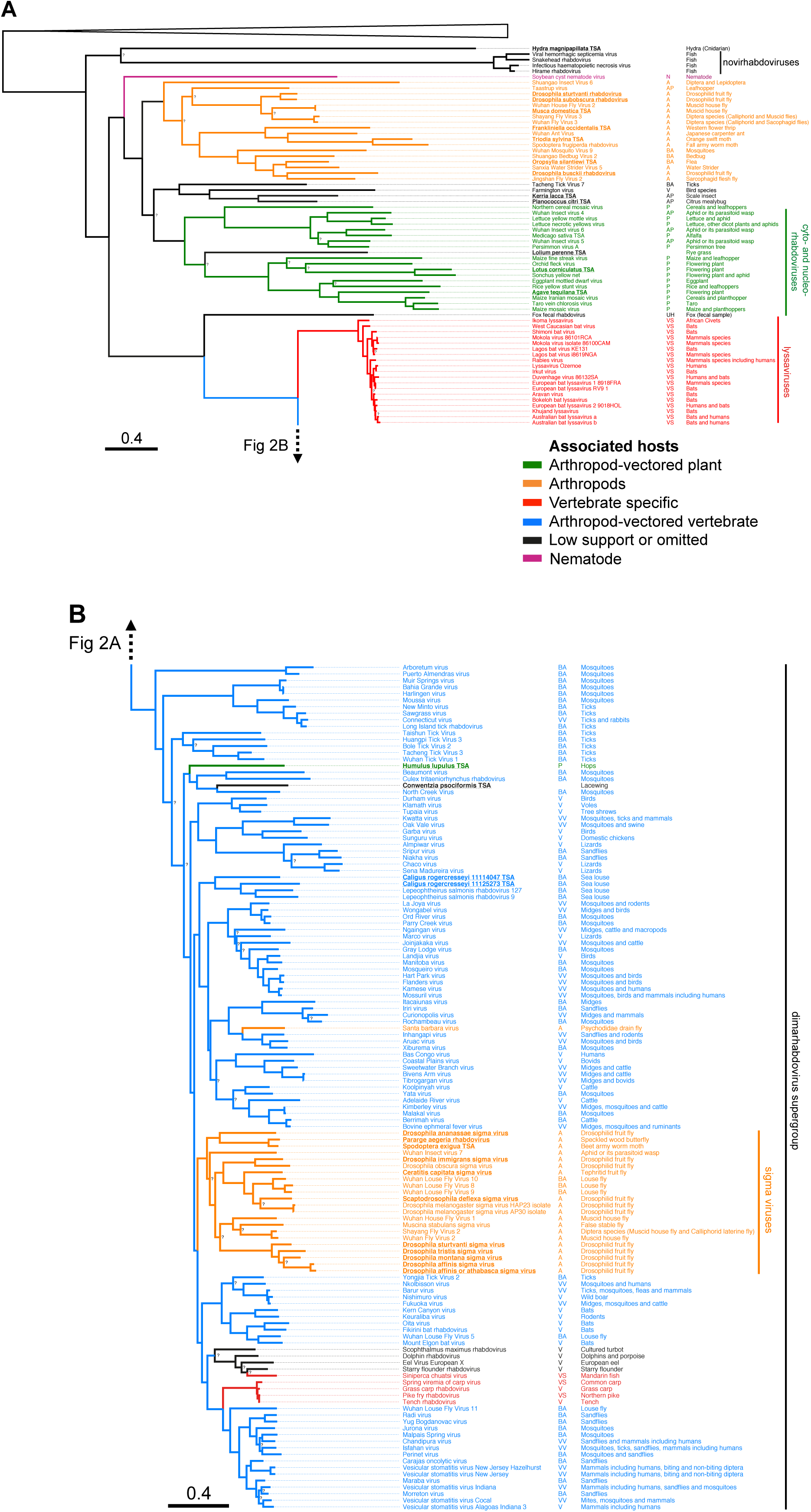
Maximum likelihood phylogeny of the *Rhabdoviridae.* (**A)** shows the basal fish-infecting novirhabdoviruses, an unassigned group of arthropod associated viruses, the plant infecting cyto- and nucleo-rhabdoviruses, as well as the vertebrate specific lyssaviruses. (**B)** shows the dimarhabdovirus supergroup, which is predominantly composed of arthropod-vectored vertebrate viruses, along with the arthropod-specific sigma virus clade. Branches are coloured based on the Bayesian host association reconstruction analysis. Black represents taxa omitted from host-state reconstruction or associations with <0.95 support. The tree was inferred from L gene sequences using the Gblocks alignment. The columns of text are the virus name, the host category used for reconstructions, and known hosts (from left to right). Codes for the host categories are: VS= vertebrate-specific, VV = arthropod-vectored vertebrate, A= arthropod specific, BS = biting-arthropod (ambiguous state), V = vertebrate (ambiguous state) AP =plant-sap-feeding-arthropod (ambiguous state), UH = uncertain-host (ambiguous across all states) and N = nematode. Names in bold and underlined are viruses discovered in this study. The tree is rooted with the Chuvirus clade (root collapsed) as identifiedas an outgroup in [13] but we note this gives the same result as midpoint and the molecular clock rooting. Nodes labelled with question marks (?) represent nodes with aLRT (approximate likelihood ratio test) statistical support values less than 0.75. Scale bar shows number of amino-acid substitutions per site. Bayesian MCC tree used to infer ancestral traits: http://dx.doi.org/10.6084/m9.figshare.1425436.

Convergence was assessed using Tracer v1.6 [48], and a burn-in of 30% was removed prior to the construction of a consensus tree, which included a description of ancestral host associations in the output file. High effective sample sizes were achieved for all parameters (>200). Previous simulations, in the context of biogeographical inference, have shown that the approach is robust to sampling bias [45]. However, to confirm this, following [49], we tested whether sample size predicts rate to or from a host association.

## Results

### Novel rhabdoviruses from RNA-seq

To search for new rhabdoviruses we collected a variety of different species of flies, screened them for CO_2_ sensitivity, which is a common symptom of infection, and sequenced total RNA of these flies by RNA-seq. We identified rhabdovirus-like sequences from a *de-novo* assembly by BLAST, and used PCR to identify which samples these sequences came from.

This approach resulted in eleven rhabdovirus-like sequences from nine (possibly ten) species of fly. Seven of these viruses were previously unknown and four had been reported previously from shorter sequences (Tables S1 and S2 http://dx.doi.org/10.6084/m9.figshare.1502665). The novel viruses were highly divergent from known viruses. Sigma viruses known from other species of *Drosophila* typically have genomes of ~12.5Kb [12, 50], and six of our sequences were approximately this size, suggesting they are near-complete genomes. None of the viruses discovered in our RNA-seq data were integrated into the host genome (see Methods for details).

To investigate the putative gene content of the viruses, we predicted genes based on open reading frames (ORFs). For the viruses with apparently complete genomes (Figure 1), we found that those from *Drosophila ananassae, Drosophila affinis, Drosophila immigrans* and *Drosophila sturtvanti* contained ORFs corresponding to the five core genes found across all rhabdoviruses, with an additional ORF between the P and M genes. This is the location of the X gene found in sigma viruses, and in three of the four novel viruses it showed BLAST sequence similarity to the X gene of sigma viruses. The virus from *Drosophila busckii* did not contain an additional ORF between the P and M genes, but instead contained an ORF between the G and L gene.

Using the phylogeny described below, we have classified our newly discovered viruses as either sigma viruses, rhabdoviruses or other viruses, and named them after the host species they were identified from (Figure 1) [51]. We also found one other novel mononegavirales-like sequence from *Drosophila unispina* that groups with a recently discovered clade of arthropod associated viruses (Nyamivirus clade [13], see Table S5 and the full phylogeny: http://dx.doi.org/10.6084/m9.figshare.1425083), as well as five other RNA viruses from various families (data not shown), confirming our approach can detect a wide range of divergent viruses.

### New rhabdoviruses from public databases

We identified a further 26 novel rhabdovirus-like sequences by searching public databases of assembled RNA-seq data with BLAST. These included 19 viruses from arthropods (Fleas, Crustacea, Lepidoptera, Diptera), one from a Cnidarian *(Hydra*) and 5 from plants (Table S3). Of these viruses, 19 had sufficient amounts of coding sequence (>1000bp) to include in the phylogenetic analysis (Table S3), whilst the remainder were too short (Table S4).

Four viruses from databases had near-complete genomes based on their size. These were from the moth *Triodia sylvina,* the house fly *Musca domestica* (99% nucleotide identity to Wuhan house fly virus 2 [13]), the butterfly *Pararge aegeria* and the medfly *Ceratitis capitata,* all of which contain ORFs corresponding to the five core rhabdovirus genes. The sequence from *C. capitata* had an additional ORF between the P and M genes with BLAST sequence similarity to the X gene in sigma viruses. There were several unusual sequences. Firstly, in the virus from *P. aegeria* there appear to be two full-length glycoprotein ORFs between the M and L genes (we confirmed by Sanger sequencing that both exist and the stop codon between the two genes was not an error). Secondly, the *Agave tequilana* transcriptome contained a L gene ORF on a contig that was the length of a typical rhabdovirus genome but did not appear to contain typical gene content, suggesting it has very atypical genome organization, or has been misassembled, or is integrated into its host plant genome [53]. Finally, the virus from *Hydra magnipapillata* contained six predicted genes, but the L gene (RDRP) ORF was unusually long. Some of the viruses we detected may be EVEs inserted into the host genome and subsequently expressed [18]. For example, this is likely the case for the sequence from the silkworm *Bombyx mori* that we also found in the silkworm genome, and the L gene sequence from *Spodoptera exigua* that contains stop codons. Under the assumption that viruses integrated into host genomes once infected those hosts, this does not affect our conclusions below about the host range of these viruses [15-17]. We also found nine other novel mononegavirale-like sequences that group with recently discovered clades of insect viruses [13] (see Table S5 and http://dx.doi.org/10.6084/m9.figshare.1425083).

### Rhabdovirus Phylogeny

To reconstruct the evolution of the *Rhabdoviridae* we have produced the most complete phylogeny of the group to date (Figure 2). We aligned the relatively conserved L gene (RNA Dependant RNA Polymerase) from our newly discovered viruses with sequences of known rhabdoviruses to give an alignment of 195 rhabdoviruses (and 26 other mononegavirales as an outgroup). We reconstructed the phylogeny using different sequence alignments and methodologies, and these all gave qualitatively similar results with the same major clades being reconstructed (Gblocks: http://dx.doi.org/10.6084/m9.figshare.1425083, TrimAl: http://dx.doi.org/10.6084/m9.figshare.1425082 and BEAST: http://dx.doi.org/10.6084/m9.figshare.1425436). The ML and Bayesian relaxed clock phylogenies were very similar: 149/188 nodes are found in both reconstructions and only 2 nodes present in the Bayesian relaxed clock tree with strong support are absent from the ML tree with strong support. These are found in a single basal clade of divergent but uniformly arthropod-specific strains, where the difference in topology will have no consequence for inference of host association. This suggests that our analysis is robust to the assumptions of a relaxed molecular clock. The branching order between the clades in the dimarhabdovirus supergroup was generally poorly supported and differed between the methods and alignments. Eight sequences that we discovered were not included in this analysis as they were considered too short, but their closest BLAST hits are listed in Table S4 (http://dx.doi.org/10.6084/m9.figshare.1502665).

We recovered all of the major clades described previously (Figure 2), and found that the majority of known rhabdoviruses belong to the dimarhabdovirus clade (Figure 2b). The RNA-seq viruses from *Drosophila* fall into either the sigma virus clade (Figure 2b) or the arthropod clade sister to the cyto- and nucleo-rhabdoviruses (Figure 2a). The viruses from sequence databases are diverse, coming from almost all of the major clades with the exception of the lyssaviruses.

### Predicted host associations of viruses

With a few exceptions, rhabdoviruses are either arthropod-vectored viruses of plants or vertebrates, or are vertebrate- or arthropod-specific. In many cases the only information about a virus is the host from which it was isolated. Therefore, *a priori,* it is not clear whether viruses isolated from vertebrates are vertebrate-specific or arthropod-vectored, or whether viruses isolated from biting arthropods (e.g. mosquitoes, sandflies, ticks, midges and sea lice) are arthropod specific or also infect vertebrates. Likewise, it is not clear whether viruses isolated from sap-sucking insects (all Hemiptera: aphids, leafhoppers, scale insect and mealybugs) are arthropod-specific or arthropod-vectored plant viruses. By combining data on the ambiguous and known host associations with phylogenetic information, we were able to predict both the ancestral and present host associations of these viruses (http://dx.doi.org/10.6084/m9.figshare.1425436). To do this we used a Bayesian phylogenetic analysis that simultaneously estimated the phylogeny and host association of our data. In the analysis we defined our host associations either as vertebrate-specific, arthropod-specific, arthropod-vectored vertebrate, arthropod-vectored plant, nematode, or as ambiguous between two (and in one case all five) of these states (see Methods).

This approach identified a large number of viruses that are likely to be new arthropod-vectored vertebrate viruses (Figure 2b). Of 80 viruses with ambiguous 89 host associations were assigned a host association with strong posterior support (>0.95). Of the 52 viruses found in biting arthropods, 45 were predicted to be arthropod-vectored vertebrate viruses, and 6 to be arthropod-specific. Of the 30 viruses found in vertebrates, 22 were predicted to be arthropod-vectored vertebrate viruses, and 2 were predicted to be vertebrate-specific (both fish viruses). Of the 7 viruses found in plant-sap-feeding arthropods (Figure 2a), 3 were predicted to be plant-associated and 2 arthropod-associated.

To test the accuracy of our predictions of current host associations we randomly selected a set of viruses with known associations, re-assigned their host association as ambiguous between all possible states (a greater level of uncertainty than we generally attributed to viruses in our data), and re-ran our analysis. We repeated this 10 times for 9 sets of 10 viruses and one set of 9 viruses (randomly sampling without replacement from the 99 viruses in our data with known host associations). These analyses correctly returned the true host association for 95/99 viruses with strong posterior support (>0.9) and 1 with weak support (mean support = 0.99, range = 0.73-1.00; Data S3 http://dx.doi.org/10.6084/m9.figshare.1538584). All three cases in which the reconstruction returned a false host association involved anomalous sequences (e.g., a change in host association on a terminal branch). Note, there would be no failure in cases where there was no phylogenetic clustering of host associations. In such cases the method would – correctly – report high levels of uncertainty in all reconstructed states.

We checked for evidence of sampling bias in our data by testing whether sample size predicts rate to or from a host association [49]. We found there is a high level of uncertainty around all rate estimates, but that there is no pattern of increased rate to or from states that are more frequently sampled.

### Ancestral host associations and host-switches

Viral sequences from arthropods, vertebrates and plants form distinct clusters in the phylogeny (Figure 2). To quantify this genetic structure we calculated the *F_st_* statistic between the sequences of viruses from different groups of hosts. There is strong evidence of genetic differentiation between the sequences from arthropods, plants and vertebrates (*P*<0.001, Figure S1 http://dx.doi.org/10.6084/m9.figshare.1495351). Similarly, viruses isolated from the same host group tend to cluster together on the tree (GSI analysis permutation tests: arthropod hosts GSI = 0.43, *P*<0.001, plant hosts GSI = 0. 46, *P*<0.001, vertebrate hosts GSI = 0.46, *P* < 0.001).

Our Bayesian analysis allowed us to infer the ancestral host association of 176 of 188 of the internal nodes on the phylogenetic tree (support >0.95), however we could not infer the host association of the root of the phylogeny, or some of the more basal nodes. A striking pattern that emerged is that switches between major groups of hosts have occurred rarely during the evolution of the rhabdoviruses (Figure 2). There are a few rare transitions on terminal branches (Santa Barbara virus and the virus identified from the plant *Humulus lupulus)* but these could represent errors in the host assignment (e.g. cross-species contamination) as well as recent host shifts. Our analysis allows us to estimate the number of times the viruses have switched between major host groups across the phylogeny, while accounting for uncertainty about ancestral states, the tree topology and root. We found strong evidence of only two types of host-switch across our phylogeny: two transitions from being an arthropod-vectored vertebrate virus to being arthropod specific (modal estimate = 2, median = 3.1, CI’s= 1.9–5.4) and three transitions from being an arthropod-vectored vertebrate virus to a vertebrate-specific virus (modal estimate = 3, median = 3.1, CI’s= 2.9–5.2). We could not determine the direction of the host shifts into the other host groups.

Vertebrate-specific viruses have arisen once in the lyssaviruses clade [3], as well as at least once in fish dimarhabdoviruses (in one of the fish-infecting clades it is unclear if it is vertebrate-specific or vector-borne from our reconstructions). There has also likely been a single transition to being arthropod-vectored vertebrate viruses in the dimarhabodovirus clade.

Insect-vectored plant viruses in our dataset have arisen once in the cyto- and nucleo-rhabdoviruses, although the ancestral state of these viruses is uncertain. A single virus identified from the hop plant *Humulus lupulus* appears to fall within the dimarhabdovirus clade. However, this may be because the plant was contaminated with insect matter, as the same RNA-seq dataset contains *COI* sequences with high similarity to thrips.

There are two large clades of arthropod-specific viruses. The first is a sister group to the large clade of plant viruses. This novel group of predominantly insect-associated viruses are associated with a broad range of insects, including flies, butterflies, moths, ants, thrips, bedbugs, fleas, mosquitoes, water striders and leafhoppers. The mode of transmission and biology of these viruses is yet to be examined. The second clade of insect-associated viruses is the sigma virus clade [11, 12, 19, 50]. These are derived from vector-borne dimarhabdoviruses that have lost their vertebrate host and become vertically transmitted viruses of insects [10]. They are common in Drosophilidae, and our results suggest that they may be widespread throughout the Diptera, with occurrences in the Tephritid fruit fly *Ceratitis capitata,* the stable fly *Muscina stabulans,* several divergent viruses in the housefly *Musca domestica* and louse flies removed from the skin of bats. For the first time we have found sigma-like viruses outside of the Diptera, with two Lepidoptera associated viruses and a virus from an aphid/parasitoid wasp. All of the sigma viruses characterised to date have been vertically transmitted [10], but some of the recently described viruses may be transmitted horizontally – it has been speculated that the viruses from louse flies may infect bats [54] and Shayang Fly Virus 2 has been reported in two fly species [13] (although contamination could also explain this result). Drosophila sigma virus genomes are characterised by an additional X gene between the P and M genes [50]. Interestingly the two louse fly viruses with complete genomes, Wuhan insect virus 7 from an aphid/parasitoid and *Pararge aegeria* rhabdovirus do not have a putative X gene. The first sigma virus was discovered in *Drosophila melanogaster* in 1937 [55]. In the last few years related sigma viruses have been found in other *Drosophila* species and a Muscid fly [10-12, 50] and here we have found sigma-like viruses in a diverse array of Diptera species, as well as other insect orders. Overall, our results suggest sigma-like viruses may be associated with a wide array of insect species.

Within the arthropod-associated viruses (the most sampled host group) it is common to find closely related viruses in closely related hosts (Figure 1). Viruses isolated from the same arthropod orders tended to cluster together on the tree (GSI analysis permutation tests: Diptera GSI = 0. 57, *P*<0.001, Hemiptera GSI = 0.34, *P*<0.001, Ixodida GSI = 0.38, *P*<0.001, Lepidoptera GSI = 0.15, *P*=0.089). This is also reflected in a positive correlation between the evolutionary distance between the viruses and the evolutionary distance between their arthropod hosts (Pearson’s correlation=0.36, 95% CI’s=0.34-0.38, *P*<0.001 based on permutation, Figure 3 and Figure S2 http://dx.doi.org/10.6084/m9.figshare.1495351). Since the virus phylogeny is incongruent with that of the respective hosts, this suggests rhabdoviruses preferentially host shift between closely related species [56, 57].

**Figure 3.**
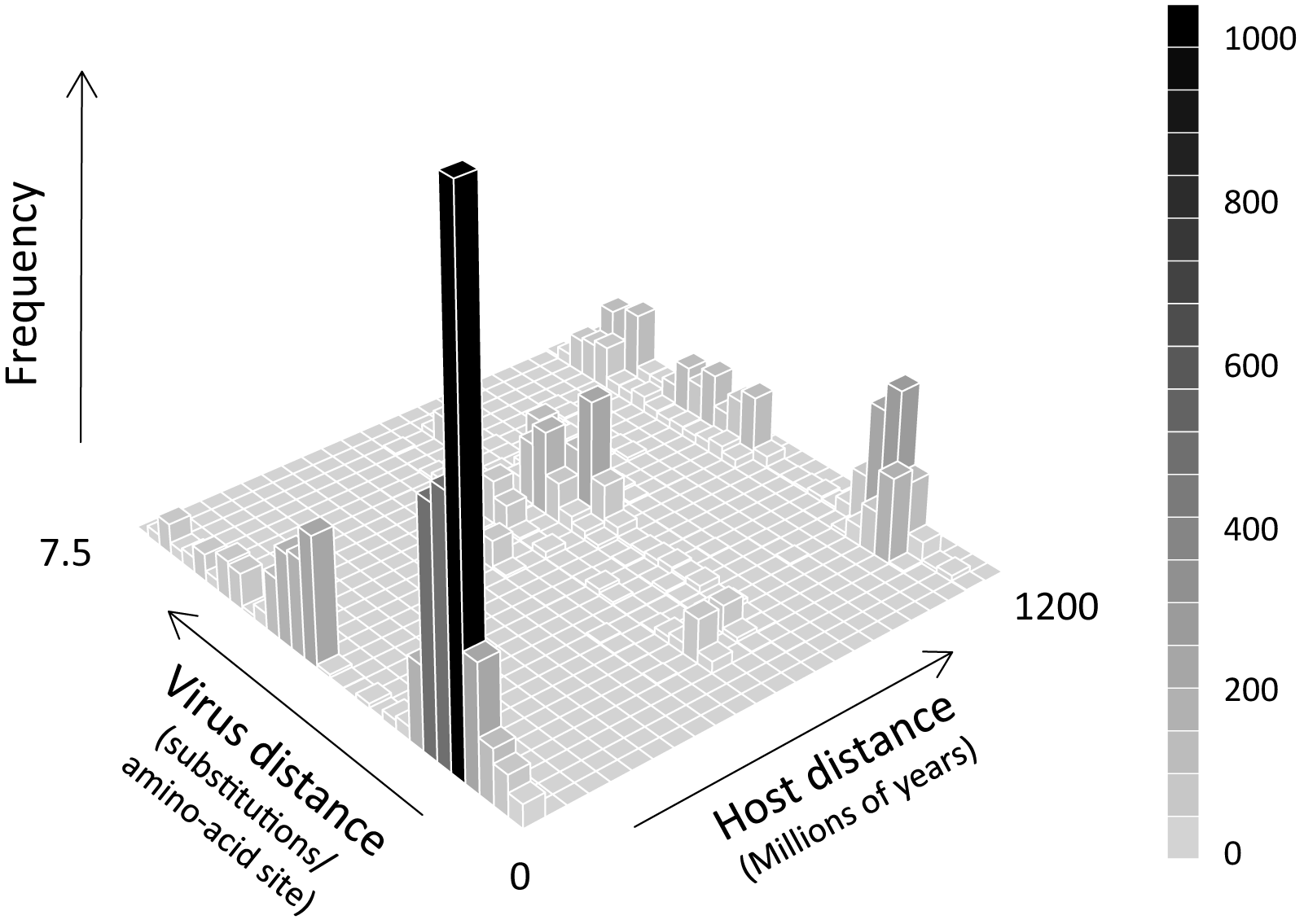
The relationship between the evolutionary distance between viruses and the evolutionary distance between their arthropod hosts (categorised by genus). Closely related viruses tend to be found in closely related hosts. Permutation tests find a significant positive correlation (correlation=0.36, 95% CI’s=0.34-0.38, *P*<0.001) between host and virus evolutionary distance (see Figure S2).

We also find viruses clustering on the phylogeny based on the ecosystem of their hosts; there is strong evidence of genetic differentiation between viruses from terrestrial and aquatic hosts (*F_st_* permutation test *P*=0.007, Figure S3 http://dx.doi.org/10.6084/m9.figshare.1495351: GSI analysis permutation tests: terrestrial hosts= 0.52. aquatic hosts= 0.29. *P*<0.001 for both). There has been one shift from terrestrial to aquatic hosts during the evolution of the basal novirhabdoviruses, which have a wide host range in fish. There have been other terrestrial to aquatic shifts in the dimarhabdoviruses: in the clades of fish and cetacean viruses and the clade of viruses from sea-lice.

## Discussion

Viruses are ubiquitous in nature and recent developments in high-throughput sequencing technology have led to the discovery and sequencing of a large number of novel viruses in arthropods [13, 14, 64]. Here we have identified 43 novel virus-like sequences from our own RNA-seq data and public sequence repositories. Of these, 32 were rhabdoviruses, and 26 were from arthropods. Using these sequences we have produced the most extensive phylogeny of the *Rhabdoviridae* to date, including a total of 195 virus sequences.

In most cases we know nothing about the biology of the viruses beyond the host they were isolated from, but our analysis provides a powerful way to predict which are vector-borne viruses and which are specific to vertebrates or arthropods. We have identified a large number of new likely vector-borne viruses – of 85 rhabdoviruses identified from vertebrates or biting insects we predict that 76 are arthropod-borne viruses of vertebrates (arboviruses). The majority of known rhabdoviruses are arboviruses, and all of these fall in a single clade known as the dimarhabdoviruses. In addition to the arboviruses, we also identified two clades of likely insect-specific viruses associated with a wide range of species.

We found that shifts between distantly related hosts are rare in the rhabdoviruses, which is consistent with previous observations that both rhabdoviruses of vertebrates (rabies virus in bats) and invertebrates (sigma viruses in Drosophilidae) show a declining ability to infect hosts more distantly related to their natural host [46, 57, 58]. It is thought that sigma viruses may sometimes jump into distantly related but highly susceptible species [56, 57, 59], but our results suggest that this rarely happens between major groups such as vertebrates and arthropods. It is nonetheless surprising that arthropod-specific viruses have arisen rarely, as one might naively assume that there would be fewer constraints on vector-borne viruses losing one of their hosts. However, this would involve evolving a new transmission route among insects, and this may be an important constraint. Within the major clades, closely related viruses often infect closely related hosts (Figure 2). For example, within the dimarhabdoviruses viruses identified from mosquitoes, ticks, *Drosophila,* Muscid flies, Lepidoptera and sea-lice all tend to cluster together (Figure 2B). However, it is also clear that the virus phylogeny does not mirror the host phylogeny, and our data on the clustering of hosts across the virus phylogeny therefore suggests that viruses preferentially shift between more closely related species (Figures 3, S1 and S2) in the same environment (Figure S3).

There has been a near four-fold increase in the number of recorded rhabdovirus sequences in the last five years. In part this may be due to the falling cost of sequencing transcriptomes [60], and initiatives to sequence large numbers of insect and other arthropods [37]. The use of high-throughput sequencing technologies should reduce the likelihood of sampling biases associated with PCR, where people look for similar viruses in related hosts. Therefore, the pattern of viruses forming clades based on the host taxa they infect is likely to be robust. However, sampling is biased towards arthropods, and it is possible that there may be a great undiscovered diversity of rhabdoviruses in other organisms [61].

Our conclusions are likely to be robust to biases in the data or limitations in the analysis. By reconstructing host associations using the Bayesian methods in the BEAST software [42] we have avoided most of the simplifying assumptions of earlier methods (e.g. symmetric transition rate matrices, lack of uncertainty associated with estimates). Nonetheless all such methods depend on there being some of sort of "process homogeneity" over the phylogeny [62]. Such analyses are of course limited by sampling; for example, if a past host is now extinct, it will never be reconstructed as an ancestral state. Nevertheless, previous studies have shown that the method is relatively robust to uneven sampling across hosts [45]. Furthermore, when we have viruses from under-sampled groups like cnidarians, fungi, nematodes, they fall outside the main clades of viruses that we are analysing. The limitations of the approach are evident in our results: we were unable to reconstruct the host associations of the root or most basal nodes of the phylogeny. The reconstructions were, however, very successful within clades that were strongly associated with a single host or clades where the less common hosts tend to form distinct subclades. As a result of this high level of phylogenetic structure, our approach was able to reconstruct the current host associations of many viruses for which we had incomplete knowledge of their host range. To check that this approach is reliable, we repeated the analysis on datasets where we deleted the information about which hosts well-characterised viruses infect. Our analysis was found to be robust, with 97% of reconstructions being accurate. The method only failed for strains with irregular host associations for their location in the phylogeny (i.e. recent changes in host on terminal branches)– a limitation that would be expected for such an analysis.

Rhabdoviruses infect a diverse range of host species, including a large number of arthropods. Our search has unearthed a large number of novel rhabdovirus genomes, suggesting that we are only just beginning to uncover the diversity of these viruses. The host associations of these viruses have been highly conserved across their evolutionary history, which provides a powerful tool to identify previously unknown arboviruses. The large number of viruses being discovered through metagenomic studies [13, 63, 64] means that in the future we will be faced by an increasingly large number of viral sequences with little knowledge of the biology of the virus. Our phylogenetic approach could be extended to predict key biological traits in other groups of pathogens where our knowledge is incomplete. However, there are limitations to this method, and the rapid evolution of RNA viruses may mean that some traits change too quickly to accurately infer traits. Therefore, such an approach should complement, and not replace, examining the basic biology of novel viruses.

## Acknowledgments

Many thanks to Mike Ritchie for providing the DMonSV infected fly line; Casper Breuker and Melanie Gibbs for PAegRV samples and Philip Leftwich for CCapSV samples. Thanks to everyone who provided fly collections. Thanks to two reviewers and Oliver Pybus for useful comments.

## Contributions

BL and FMJ conceived and designed the study. BL and JD carried out molecular work. BL, WJP, DJP and DJO carried out bioinformatic analysis. BL, GGRM and JJW carried out phylogenetic analysis. BL GGRM, JJW and FMJ wrote the manuscript with comments from all other authors. All authors gave final approval for publication.

## Funding

BL and FMJ are supported by a NERC grant (NE/L004232/1), a European Research Council grant (281668, DrosophilaInfection), a Junior Research Fellowship from Christ’s College, Cambridge (BL). GGRM is supported by an MRC studentship. The metagenomic sequencing of viruses from *D. immigrans, D. tristis* and *S. deflexa* was supported by a Wellcome Trust fellowship (WT085064) to DJO.

## Supplementary materials

Tables S1-5. List of newly discovered viruses:

Figures S1-S3

Supplementary Figure 1.

Results of permutation tests of population differentiation between the sequences of viruses taken from different categories of host: (a) between arthropods and vertebrates, (b) between arthropods and plants and (c) between plants and vertebrates. Red lines show the degree of differentiation, measured using Hudson’s *F_st_* estimator, and grey histograms show the *F_st_* values from 1000 unique random permutations of host categories over viral sequences. We found significant differentiation between each pair of categories (*F_st_* values: a=0.07, b=0.35, c=0.48) with none of the 1000 permutations resulting in as high an *F_st_* value as the data (*P*<0.001 for all).

Supplementary Figure 2.

Correlation of the evolutionary distances between rhabdovirueses and the evolutionary distances between their arthropod hosts. Figure shows the results of a permutation test of this relationship (measured by Pearson’s correlation coefficient). The red line shows the true correlation coefficient (0.36), and the grey histogram shows the correlation coefficients returned from 1000 unique random permutations of host genera over viral sequences (see Methods). We found a significant association between viral and host phylogenies with none of the 1000 permutations resulting in as strong a correlation as the data (*P*<0.001).

Supplementary Figure 3.

Results of the permutation test of population differentiation between the sequences of viruses taken from land and aquatic arthropod and vertebrate hosts. Only arthropod and vertebrate hosts were included in the test, since aquatic hosts only included arthropods and vertebrates. The red line shows the true degree of differentiation, measured using Hudson’s *F_st_* estimator, and the grey histogram shows the *F_st_* values returned from 1000 unique random permutations of host categories over viral sequences. Viruses from terrestrial and aquatic hosts were found to have significant population differentiation (with only 7 of the 1000 permutations resulted in as high an *F_st_* value as the data *(F_st_* =0.07, *P*=0.007).

## All data has been made available in public repositories

NCBI Sequence Read Archive Data: SRP057824

Data S1, sample information: http://dx.doi.org/10.6084/m9.figshare.1425432

Data S2, virus ID, Genbank accession numbers and host information: http://dx.doi.org/10.6084/m9.figshare.1425419

Data S3, results from testing ancestral trait reconstructions predictions: http://dx.doi.org/10.6084/m9.figshare.1538584

L gene sequences fasta: http://dx.doi.org/10.6084/m9.figshare.1425067

TrimAl alignment fasta: http://dx.doi.org/10.6084/m9.figshare.1425069

Gblocks alignment fasta: http://dx.doi.org/10.6084/m9.figshare.1425068

Phylogenetic tree Gblocks alignment: http://dx.doi.org/10.6084/m9.figshare.1425083

Phylogenetic tree TrimAl alignment: http://dx.doi.org/10.6084/m9.figshare.1425082

BEAST alignment fasta: http://dx.doi.org/10.6084/m9.figshare.1425431

BEAUti xml file: http://dx.doi.org/10.6084/m9.figshare.1431922

Bayesian analysis tree: http://dx.doi.org/10.6084/m9.figshare.1425436

